# Excess light priming in *Arabidopsis thaliana* genotypes with altered DNA methylomes

**DOI:** 10.1101/475798

**Authors:** Diep R. Ganguly, Bethany A. B. Stone, Andrew F. Bowerman, Steven R. Eichten, Barry J. Pogson

**Author notes:** These authors contributed equally to this work (Diep R. Ganguly, Bethany A. B. Stone). **Corresponding author:** Barry Pogson. **Author contributions:** DRG, BABS, SRE, and BJP conceived and designed the study; DRG, BABS, and AFB designed experiments; BABS performed the experimental work with guidance from DRG and AFB; DRG and BABS performed data analyses with guidance from SRE and AFB; DRG and BABS drafted the manuscript; all authors commented on and edited the manuscript.

## Abstract

Plants must continuously react to the ever-fluctuating nature of their environment. Repeated exposure to stressful conditions can lead to priming, whereby prior encounters heighten a plant’s ability to respond to future events. A clear example of priming is provided by the model plant *Arabidopsis thaliana* (Arabidopsis), in which photosynthetic and photoprotective responses are enhanced following recurring light stress. While there are various post-translational mechanisms underpinning photoprotection, an unresolved question is the relative importance of transcriptional changes towards stress priming and, consequently, the potential contribution from DNA methylation – a heritable chemical modification of DNA capable of influencing gene expression. Here, we systematically investigate the potential molecular underpinnings of physiological priming against recurring excess-light (EL), specifically DNA methylation and transcriptional regulation: the latter having not been examined with respect to EL priming. The capacity for physiological priming of photosynthetic and photoprotective parameters following a recurring EL treatment was not impaired in Arabidopsis mutants with perturbed establishment, maintenance, or removal of DNA methylation. Importantly, no differences in development or basal photoprotective capacity were identified in the mutants that may confound the above result. Little evidence for a causal transcriptional component of physiological priming was identified; in fact, most alterations in primed plants presented as a transcriptional ‘dampening’ in response to an additional EL exposure, likely a consequence of physiological priming. However, a set of transcripts uniquely regulated in primed plants provide preliminary evidence for a novel transcriptional component of recurring EL priming, independent of physiological changes. Thus, we propose that physiological priming of recurring EL in Arabidopsis occurs independently of DNA methylation; and that the majority of the associated transcriptional alterations are a consequence, not cause, of this physiological priming.

**One sentence summary:** Photoprotection and priming against recurring excess light is functional despite impaired maintenance of the DNA methylome.

## Introduction

Plants must respond to various stresses imposed by their environment. Abiotic stresses may occur over long periods of development with unfavourable conditions persisting stably as is characteristic of extreme climates. Alternatively, environments can be highly dynamic and comprise of transient, often recurring, stressful events. Information processing is key for effective physiological and developmental responses to specific environmental factors. Indeed, there is growing evidence that plants can ‘remember’ past experiences (Hilker *et al.* 2016). In addition to long-term acclimation to sustained environmental changes, short-term plant stress responses are modified by prior exposure to a transient, and often recurring, specific environmental stimulus (referred to as priming). Here, the future fitness of a primed individual is increased by reducing the damage of stressful events, while the costs of initiating and maintaining priming are outweighed by the costs of stress exposure in an ‘un-primed’ (or naive) state (Hilker *et al.* 2016).

A variety of mechanisms have been reported to contribute towards stress priming including transcriptional memory underpinned by stalled RNA Pol II and elevated H3K4me3 (Ding *et al.* 2012), and fractionation of H3K27me3 patterns (Sani *et al.* 2013). It has also been reported that the activity of the HSFA2 transcription factor can result in H3K4me2 and H3K4me3 changes, in response to recurring heat stress, to convey transcriptional priming (Lämke *et al.* 2016). Additionally, HDA6-mediated histone H4 de-acetylation has been linked to prime jasmonic acid signalling leading to enhanced drought tolerance (Kim *et al.* 2017). These various examples highlight the importance of chromatin variation in facilitating of plant stress priming. Another chromatin mark speculated to promote priming is DNA methylation, variations in which could, theoretically, be stably inherited over mitotic cell divisions to convey persistent transcriptional control (Johannes and Schmitz 2018). The targeting of the RNA-directed DNA methylation (RdDM) pathway towards promoter regions of genes, and the occurrence of gene body methylation (gbM), suggests that differential methylation could arise within genes or their regulatory elements to cause functional differences (Matzke and Mosher 2014; Bewick *et al.* 2016). Such observations highlight the potential regulatory capacity for DNA methylation.

An extant question is the exact regulatory potential of DNA methylation. Canonically, DNA methylation is considered a mechanism for transcriptional repression, for example through steric hindrance of RNA polymerase II (Molloy 1986). This silencing is most pronounced at transposable elements (TEs) (Cokus *et al.* 2008). On the other hand, gbM is often found within constitutively expressed genes although there is conflicting evidence for an effect on transcription (Bewick *et al.* 2016; Muyle and Gaut 2018). Other reports implicate the involvement of DNA methylation in alternative splicing and modulation of transcription factor binding capacity (Shukla *et al.* 2011; O’Malley *et al.* 2016; Yin *et al.* 2017). Given these various mechanisms, and placing DNA methylation in the broader context as being one of many chromatin modifications, it might be unsurprising that efforts to quantify the contribution of DNA methylation, at an organism level, towards transcription have shown a weak relationship (Meng *et al.* 2016). Instead, changes in the methylome may be a consequence of gene expression changes rather than a driver (Secco *et al.* 2015). The ability to identify causative changes in the methylome are further complicated as the effects of DNA methylation can be in both *cis* and *trans* (Rowley *et al.* 2017).

Given these complications, various tools exist to quantify the effects of variable methylation (epi-alleles) on, and in response to, gene expression and physiological traits. The utilization of epigenetic recombinant inbred lines that display variable methylation patterns, but are isogenic, demonstrated that epi-alleles could contribute towards quantifiable phenotypic differences (Johannes *et al.* 2009; Cortijo *et al.* 2014). Mutants with defective methylation machinery that exhibit a variety of methylome variations, depending on the severity of the mutation, have also been used to correlate a relationship between DNA methylation and plant stress responses (Boyko *et al.* 2010; Le *et al.* 2014; Wibowo *et al.* 2016). Mutants are also described to be developmentally or morphologically aberrant, however, it is not clear whether this is the direct result of methylation changes or an indirect effect of TE de-regulation and genomic instability (Finnegan *et al.* 1996; Reinders *et al.* 2009; Stroud *et al.* 2014; Williams and Gehring 2017). Indeed, traits attributed towards methylome variants could equally be tied to underlying TE activity (Ong-Abdullah *et al.* 2015; Wibowo *et al.* 2016; He *et al.* 2018).

We previously demonstrated that Arabidopsis is primed by a recurring EL regime, evident by altered non-photochemical quenching (NPQ) and improved photosystem II (PSII) efficiency (Ganguly *et al.* 2018). However, we detected no associated changes in DNA methylation. While this demonstrated that the Arabidopsis methylome was impervious to recurring EL, it does not preclude transcriptional regulation of EL priming to which appropriate methylome maintenance may be important. In fact, EL-exposed tissue can promote the induction of EL-responsive transcripts in naive leaves for added photoprotection through the process of systemic acquired acclimation (SAA) (Karpinski 1999; Rossel *et al.* 2007; Gordon *et al.* 2012). As the methylation machinery was operational during previous analyses, any light-induced changes may have been reset prior to tissue harvesting. Indeed, the disruption of methylation pathways has revealed transgenerational effects that were otherwise reset (Iwasaki and Paszkowski 2014). Thus, we sought to clarify these unknowns by testing whether a range of mutants, which are unable to maintain or reset their methylome, were capable of priming against recurring EL; and subsequently characterising differences in the transcriptome of EL primed plants. We report that DNA methylation mutants display functional photoprotection and EL priming to an equivalent extent as wild-type (WT; Col-0) plants. Furthermore, whilst primed plants demonstrate a completely reset transcriptome, they also displayed attenuated responses to further EL, which we refer to as “dampening”, potentially reflecting the reduced generation of stress signalling molecules due to enhanced photoprotection.

## Results

### Mutant characterisation and the novel *strs2* methylome

We utilized three Arabidopsis T-DNA insertion mutants targeting components of the DNA methylation machinery to test for stress priming with disrupted methylome maintenance. These include the triple methyltransferase mutant *drm1drm2cmt3* (*ddc3*) (Henderson and Jacobsen 2008), the triple demethylase mutant *ros1dml2dml3* (*rdd*) (Le *et al.* 2014), and the novel putative RdDM mutant *strs2* (Khan *et al.* 2014). While the *ddc3* and *rdd* mutants show gross methylome changes across the genome, *strs2* displays subtle effects on the methylome by fine-tuning DNA methylation levels at stress-associated genes (Khan *et al.* 2014). As previous studies investigating *strs2* relied on a targeted analysis using chop-PCR, we performed MethylC-seq (n=3) to confirm the extent of methylome changes. Global levels of CG, CHG, and CHH methylation in *strs2* were highly comparable to WT (Figure 1 A, Supplementary Dataset 1). However, employing *DSS* (Feng *et al.* 2014; Wu *et al.* 2015) to identify differentially methylated regions (DMRs) revealed moderate levels of local changes, predominantly in the CG context, with comparable numbers of hyper- and hypo-DMRs (Figure 1 B, Table 1, Supplementary Dataset 2). Nonetheless, many CHH DMRs were located within TEs consistent with the implication of STRS2 in RdDM. Furthermore, 130/199 (65.6%) *strs2* CHH hypo-DMRs were regions targeted by DRM1 and DRM2 (7,393 CHH hypo-DMRs in *drm1drm2*), whereas CG and CHG hypo-DMRs demonstrated weaker overlaps (CG: 7/318, 2.2%; CHG: 8/34, 23.5%). Paired with previous results this suggests that *strs2* is a “weak” RdDM mutant (Stroud *et al.* 2015).

**Table 1.** DMR analysis in *strs2* (vs WT)

| DMR type | Methylation context |  |  |  |
| --- | --- | --- | --- | --- |
|  | CG | CHG | CHH | Total |
| Hyper-methylation | 700 | 68 | 148 | 916 |
| Hypo-methylation | 684 | 38 | 226 | 948 |
| Total | 1,384 | 106 | 374 | 1,864 |

**Figure 1.**
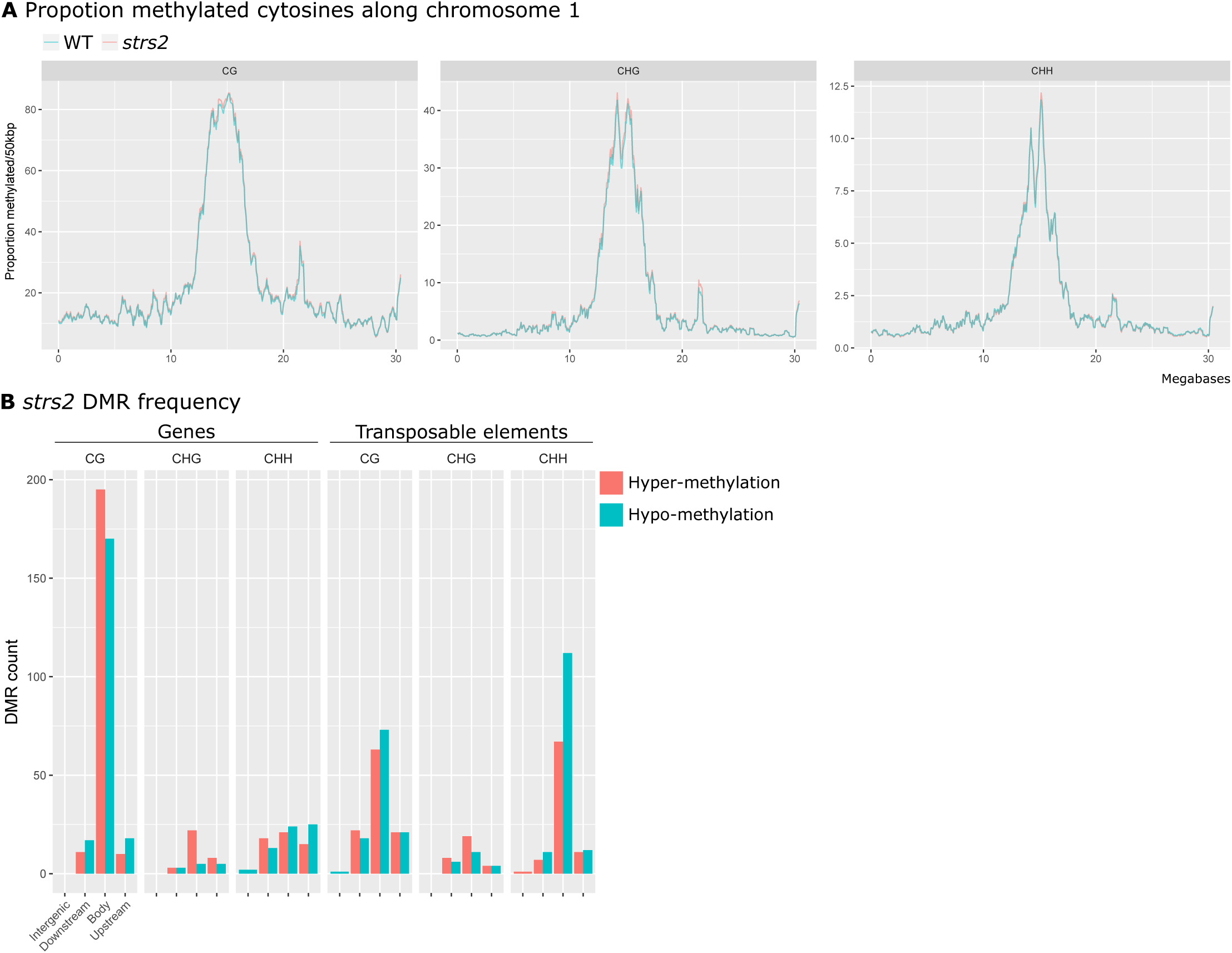
Subtle methylome perturbation in *strs2*. **A** Mean weighted methylation levels, binned into 50 kbp rolling windows, along Arabidopsis chromosome 1 for WT and *strs2*. **B** DMR frequency in *strs2* grouped by genomic location relative to annotated genes or transposable elements. DMRs were classified as either occurring directly within (body), <1 kbp away from the 5’ (upstream) or 3’end of (downstream), or > 1 kbp (intergenic) away from genomic features.

### Development and basal photoprotection in methylome mutants

Aberrant DNA methylation is considered to result in developmental differences due to transcriptional mis-regulation and TE activation; *ddc3* displays curled leaves and reduced stature due to *SDC* mis-expression (Henderson and Jacobsen 2008), while *strs2* shows a slight early flowering phenotype (Kant *et al.* 2007). To ensure that developmental abnormalities would not confound observations of priming, developmental traits were monitored until 4-weeks of age starting from 10-day old seedlings (Figure 2). While no unusual phenotypes were evident for *rdd* and *strs2* (Figure 2 A), *ddc3* displayed the expected curled leaves after 3-weeks of age. Quantitative measures of plant area reflect these visible observations, whereby the rosette area of *ddc3* was reduced compared to WT (Figure 2 B). Minor differences in leaf number and plant area were observed between 3 and 4 weeks for *rdd* and *strs2*. Therefore, priming measurements were performed on 3 week-old plants prior to the onset of substantial morphological differences.

**Figure 2.**
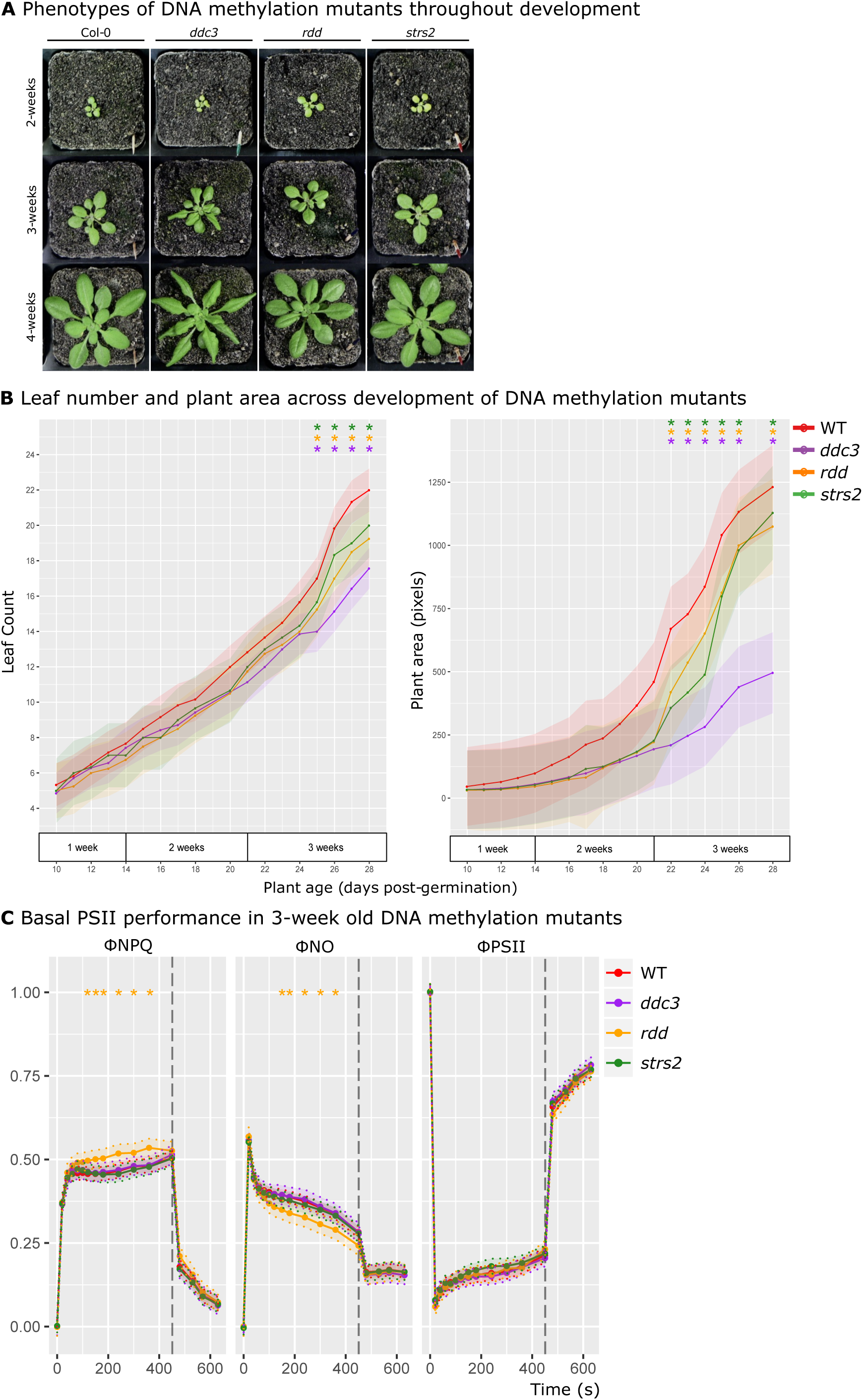
DNA methylation mutant development and basal PSII function. **A** Representative morphology of WT, *ddc3, rdd*, and *strs2* from 2-4 weeks of age. **B** Leaf number (left) and plant area (right) in 10-day old WT (red, n=8), *ddc3* (purple, n=8), *rdd* (orange, n=8), and *strs2* (green, n=8) seedlings, monitored for 20 days until 4-weeks of age. **C** Measures of PSII performance in 3-week old WT (red, n=60), *ddc3* (purple, n=10), *rdd* (orange, n=10), *strs2* (green, n=15) plants grown under standard conditions. Points denote estimated marginal means based on a fitted linear mixed-effect model for each genotype. Bars and shaded regions denote 95% confidence intervals; * indicates statistical significance (adjusted p-value<0.05) from WT.

In addition to developmental abnormalities, aberrant methylome maintenance may also affect the capacity for basal photoprotective responses. Thus, we tested functional PSII capacity in 3-week old mutants (Figure 2 C). Photosynthetic efficiency (ΦPSII) was largely consistent across genotypes. Similarly, the capacity for, or activation of, actively regulated (ΦNPQ) and constitutively (ΦNO) dissipative quenching was largely consistent across genotypes. An exception to this was *rdd* that demonstrated elevated ΦNPQ with a concomitant reduction in ΦNO but unperturbed ΦPSII. However, the difference observed in *rdd* is minor compared to traditional mutants with perturbed photosystems (Niyogi *et al.* 2001) and is more reflective of natural variation (Jung and Niyogi 2009). Thus, no major disruptions to basal PSII performance is evident in these methylation mutants allowing for comparisons of EL priming.

### Excess light priming evident despite aberrant methylome patterning

To investigate whether a perturbed methylome would impair priming to recurring EL we repeated the WLRS time-course experiment (Ganguly *et al.* 2018) on WT, *ddc3, rdd*, and *strs2* (Figure 3). The week of recurring EL led to the expected morphological differences across all genotypes, namely increased rosette compaction but consistent leaf area (Figure 3 A-B). However, there was an attenuated difference in *ddc3* area, which we attribute to its naturally curled leaf phenotype. Alongside morphological measures were changes in PSII traits indicative of physiological priming to recurring EL (Figure 3 C). The adoption of ΦNPQ and ΦNO also revealed distinct patterns in thermal dissipation not observed previously. Particularly, that the enhanced NPQ was underpinned by actively regulated pH-dependent thermal dissipation (ΦNPQ) rather than due to constitutive forms (ΦNO). Indeed, ΦNO was lower in primed plants. While ΦNPQ is rapidly activated in primed plants, it is deactivated concomitantly with an increase in ΦPSII. A key result here was that all methylation mutants demonstrated a WT-like priming response, for all three PSII traits, to recurring EL thus demonstrating appropriate light-acclimatory processes despite having perturbed methylomes.

**Figure 3.**
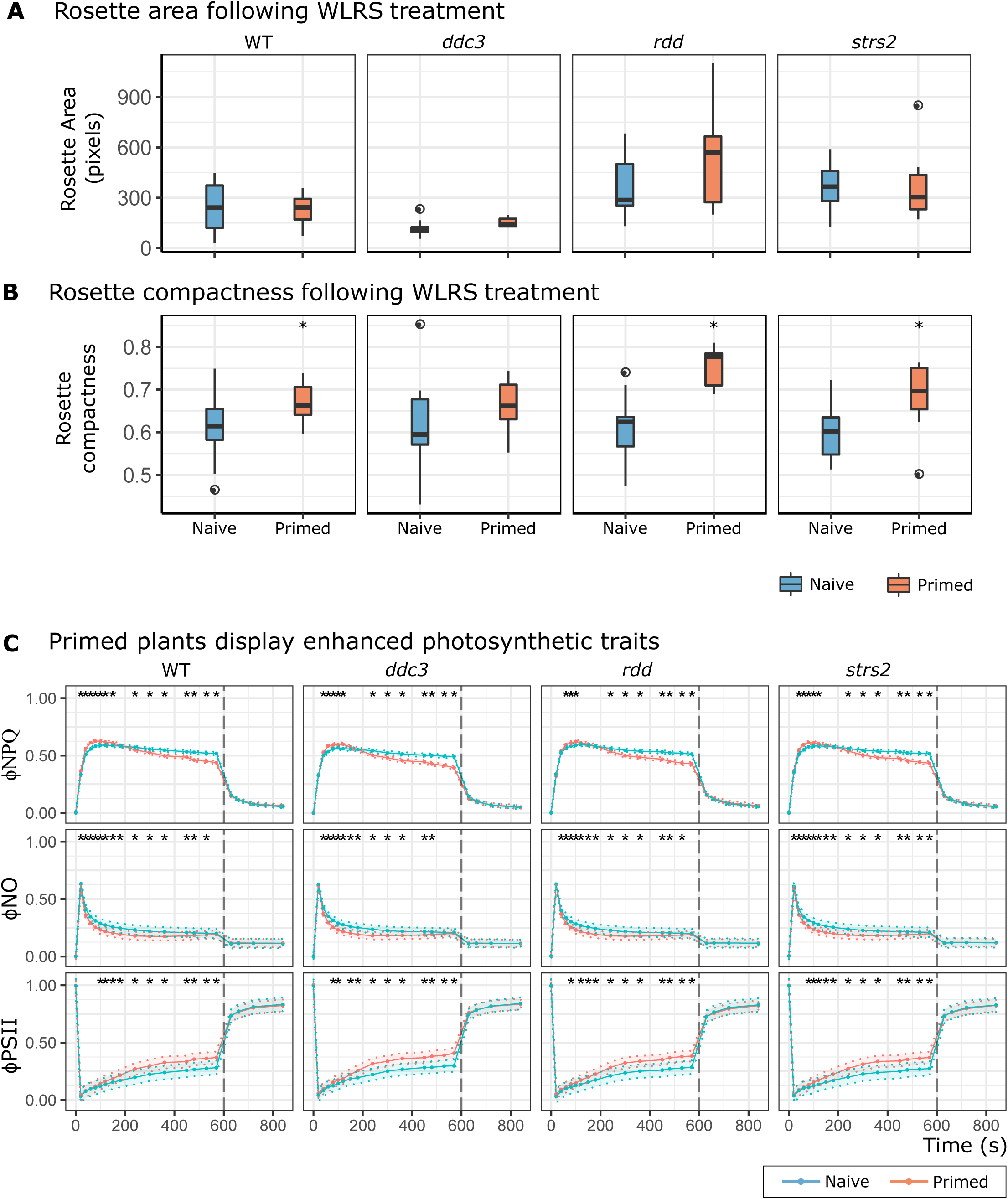
DNA methylation mutants exhibit priming to recurring EL. **A-B** Boxplots of rosette area and compactness measured in all genotypes either exposed to recurring EL (primed, n=12-15) or not (naive, 21-23). * denotes statistical significance determined using independent Student’s t-tests for each genotype (adjusted p-value < 0.05). **C** PSII performance traits in naive (WT n=22; *ddc3* n=16, *rdd* n=21, *strs2* n=23) and primed (WT n=16; *ddc3* n=13, *rdd* n=12, *strs2* n=15) plants for all genotypes. Points denote estimated marginal means based on a fitted linear mixed-effect model for each genotype. Bars and shading denote 95% confidence intervals; * indicates statistical significance (adjusted p-value < 0.05) from WT.

### Primed plants exhibit transcriptional ‘dampening’ despite a reset transcriptome

Persistent changes in gene expression are hypothesized to convey a primed state (Hilker *et al.* 2016). To test this, we searched for constitutive changes in gene expression between naïve (N) and primed (P) plants (n=3) after one-week of recurring EL using mRNA-seq. Despite the observation of physiological priming in these tissues, we observed a completely reset transcriptome exemplified by limited variance and 0 differentially expressed genes (DEGs) between naïve and primed samples (Figure S1 A-B). We subsequently investigated the transcriptional response of primed plants to an additional EL treatment (triggering stress, P+T). Application of an EL triggering stress to naive plants (N+T) elicited the differential expression of hundreds of genes, however, primed plants showed a vastly attenuated response (Figure S1 C-D, Supplementary Datasets 4 - 5). While the majority of EL-induced transcripts in P+T plants overlapped with those in N+T plants, a striking proportion of both up- (73%) and down-regulated (85%) transcripts were unique to N+T plants (Figure 4 A). The abundance of nearly all 318 uniquely down-regulated transcripts in N+T plants was greater in P+T plants (Figure 4 B, left). Equivalently, abundance of the 256 uniquely up-regulated transcripts in N+T plants was decreased in P+T plants (Figure 4 B, right). Together, these results evoke a dampened transcriptional response to EL triggering stimuli in primed plants, rather than reflecting differences between their basal (un-triggered) transcriptomes. The dampening effect is, however, subtle in nature - no DEGs were identified upon direct comparison of N+T and P+T samples.

**Figure 4.**
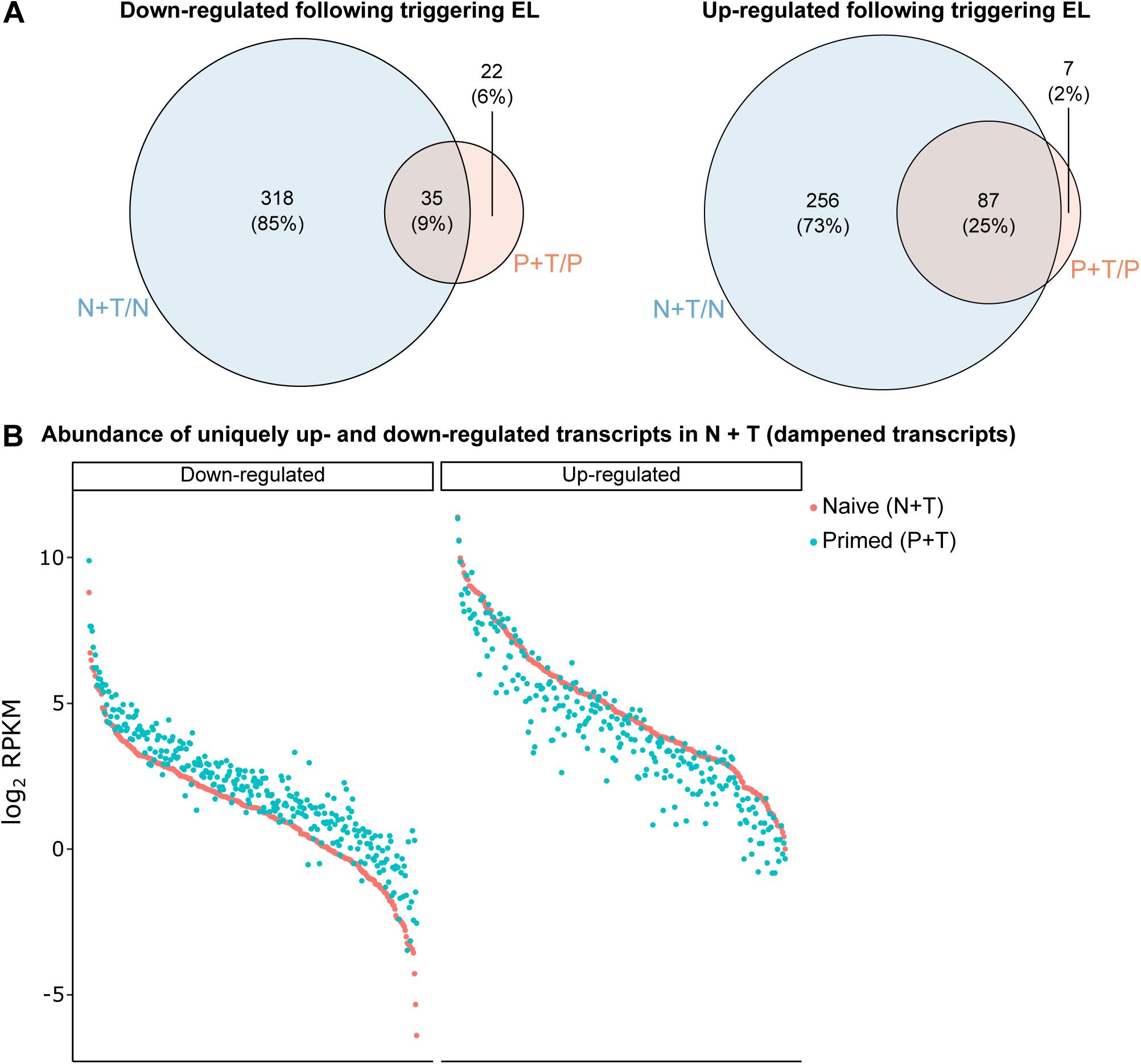
Transcriptional ‘dampening’ evident in EL primed plants. **A** Overlap between up- and down-regulated transcripts in naive and primed plants in response to an additional triggering EL treatment. **B** Transcript abundance (log_2_ RPKM) of up-(left) and down-regulated (right) transcripts, in N+T plants, for both N+T and P+T plants. **C** Identifying dampened transcripts that are inducible by various stress signals. * denotes significant overlaps (p<0.05) as determined by a hypergeometric test using *phyper*.

We hypothesised that the transcriptional dampening observed in primed plants may be a consequence of reduced signalling because of enhanced dissipative quenching (Rossel *et al.* 2007; Suzuki *et al.* 2013; Carmody *et al.* 2016). Indeed, a gene ontology analysis of all 574 transcripts exhibiting dampening (Supplementary Dataset 6), in primed plants, revealed an enrichment for terms associated with response to abiotic stimuli, including light intensity, as well as response to ROS and hydrogen peroxide, supporting this hypothesis (Supplementary Dataset 7). Genes induced by various ROS (hydrogen peroxide [H_2_O_2_], singlet oxygen [^1^O_2_], and superoxide [O^2-^] (Gadjev *et al.* 2006)), salicylic acid (SA) (Blanco *et al.* 2009), β-cyclocitral (Ramel *et al.* 2012), ABA (Pornsiriwong *et al.* 2017), and SAA (Rossel *et al.* 2007) were overlapped with the dampened transcripts identified here. In total, 198/574 (34.5%, P_[X≥198]_ < 0.05) dampened transcripts were overlapped from the collated gene list. This was driven by significant overlaps with ABA, β-Cyclocitral, and SAA induced genes as determined by hypergeometric testing (Table 2).

**Table 2.** Hypergeometric testing of 574 dampened transcripts overlapped with genes involved in stress signalling

| Gene list | Total responsive genes | Overlap with dampened transcripts | Hypergeometric test |
| --- | --- | --- | --- |
| ABA | 2,481 | 104 | $P_{[X \geq 104]} < 0.01$ |
| SA | 217 | 7 | $P_{[X \geq 7]} = 0.56$ |
| O<sub>2</sub><sup>-</sup> | 209 | 5 | $P_{[X \geq 5]} = 0.81$ |
| H<sub>2</sub>O<sub>2</sub> | 325 | 11 | $P_{[X \geq 11]} = 0.49$ |
| <sup>1</sup>O<sub>2</sub> | 295 | 10 | $P_{[X \geq 10]} = 0.49$ |
| β-Cyclocitral | 1,145 | 61 | $P_{[X \geq 61]} < 0.001$ |
| SAA | 703 | 41 | $P_{[X \geq 41]} < 0.001$ |

## Discussion

The popular notion that DNA methylation may regulate plant stress responses is confounded by both supporting and conflicting investigations spanning a range of species and stressors. Light stress is one such example for which physiological priming and memory have been demonstrated in Arabidopsis (Szechyńska-Hebda *et al.* 2010; Gordon *et al.* 2012), yet we did not observe co-occurring stress-induced DNA methylation changes (Ganguly *et al.* 2018). Consequently, we refined the experimental design to address a distinct hypothesis: that physiological priming of recurring EL is underpinned by transcriptional regulation for which active maintenance of DNA methylation is important. We present evidence for recurring EL priming in DNA methylation mutants using a full-factorial experimental design, which facilitated systematic testing of two closely related potential priming mechanisms: 1. DNA methylation-mediated transcriptional regulation; and 2. altered transcription (independent of DNA methylation).

### DNA methylation, development, and priming

Loss of DNA methylation leads to abnormal plant growth due to its importance for genome integrity (Reinders *et al.* 2009). Abnormal leaf development could in turn lead to confounding changes in light acclimation that are not a direct result of DNA methylation changes to light stress-responsive genes. Thus, we performed detailed physiological characterisation of DNA methylation mutants to identify any constitutive changes in photosynthetic rate or capacity that could confound our interpretation of the contribution of DNA methylation to priming. We found negligible evidence of changes to photosynthesis at the developmental stage examined here.

Light stress induces multiple physiological, post-transcriptional and extensive transcriptional responses (Dietz 2015). Thus, it was hypothesized that changes in DNA methylation might impact the transcriptional component of the acclimation response, thereby impairing the ability for EL priming. Contradictory to this hypothesis perturbation of the methylome, regardless of severity (e.g. *strs2* vs *ddc3*) or type (e.g. hyper- or hypo-methylation), did not impair basal photoprotective capacity nor the ability to be primed by recurring EL. Instead, the EL priming observed herein likely reflects post-transcriptional changes of the photosynthetic machinery. The reduction in ΦNO disputes constitutive changes in PSII conformation. Rather there appears to be greater electron flow in primed plants indicated by increased ΦPSII, which may reflect an increased concentration of components facilitating photosynthetic electron transport, such as PSII core proteins, cytochrome b_6_f, and chloroplastic ATPases (Foyer *et al.* 2012; Dwyer *et al.* 2012; Zhu *et al.* 2017). Pertinent to this study is the observation that these processes remain functional under a range of methylome perturbations.

### Priming occurs independently of transcriptional changes

Transcriptional up-regulation of light-responsive genes is indicative of an acclimatory response (Gordon *et al.* 2012; Carmody *et al.* 2016), however, it remains an untested hypothesis that EL priming is underpinned by constitutive transcriptional changes. No persistent changes in gene expression were observed in primed plants, consistent with previous reports of rapid transcriptome resetting following EL treatment (Crisp *et al.* 2017). An attenuated transcriptional response (“transcriptional dampening”) was instead observed in primed plants responding to a final triggering stress. This contrasts to the enhanced transcriptional responses previously observed upon a triggering stimulus following single abiotic stress treatments (Ding *et al.* 2012; Lämke *et al.* 2016; Crisp *et al.* 2017). Instead, an attenuated transcriptional response likely reflects a greater capacity of primed plants, with longer-term recurring EL exposure, to deal with increased light flux. That is, enhanced photosynthetic and photoprotective capacity in primed plants may alter the extent of EL-mediated retrograde signalling, leading to an attenuated response. Indeed, changes in the rate of electron flow through the photosynthetic apparatus are known to alter the modulation of nuclear gene expression. For instance, chloroplast regulation of nuclear gene expression under EL was impaired by disrupting effective photosynthetic electron flow causing PSI photodamage and attenuating oxylipin biosynthesis and signalling (Gollan *et al.* 2017). Additionally, impairing photosynthetic electron flow, thus altering the redox poise of the plastoquinone pool, attenuates the transcriptional responses required for thermal acclimation (Dickinson *et al.* 2018). Taken together, we propose that the dampened transcriptional response might be a consequence of altered retrograde signalling due to physiological priming. In support of this hypothesis was the observation that approximately one-third of dampened transcripts overlapped with genes induced by signalling pathways, especially ABA, β-Cyclocitral, and SAA. These results suggest that the gene expression changes observed in primed plants are consequential of physiological priming, rather than the cause.

## Conclusion

Through the analysis of Arabidopsis mutants impaired in active methylome maintenance, we have conducted a systematic investigation of the hypothesis that DNA methylation may contribute to stress priming. Specifically, we address the hypothesis that physiological priming to recurring EL is underpinned by transcriptional regulation and active methylome maintenance. We found no evidence to suggest that aberrant maintenance or removal of DNA methylation impaired the capacity for physiological priming against recurring EL. In fact, primed plants demonstrated completely reset transcriptomes. Instead, an attenuation in transcriptional response to EL was observed in primed plants and is likely to be a consequence rather than a cause of physiological priming.

## Methods

### Plant growth and germplasm

All *Arabidopsis* germplasm utilized were in the Columbia (Col-0) background. Plant lines comprised of wild-type Col-0 (WT), the DNA methylation establishment triple mutant *ddc3* (CS16384; *drm1*-2 *drm2*-2 *cmt3*-11), the “weak” RdDM mutant *strs2* (SALK_028850), and the triple demethylase mutant *rdd* (derived from *ros1-4*, SALK_045303; *dml2*, SALK_131712; *dml3-2*, SALK_056440). All seeds were obtained from the Arabidopsis Biological Research Centre (Ohio State University, Columbus, OH, USA), with the exception of *rdd* which was kindly provided by Dr. Ming-Bo Wang (CSIRO, Canberra, Australia) (Le *et al.* 2014). Primers used for genotyping are listed in Supplementary Dataset 8.

For plant growth, Arabidopsis seeds were sown onto individual pots containing moist Seed Raising Mix (Debco, NSW, Australia). Soil was supplemented with Osmocote Exact Mini slow release fertilizer (Scotts, NSW, Australia) at a concentration of 1 g/L dry volume of soil and treated with 1 L of 0.3 % (v/v) AzaMax (OCP, NSW, Australia) prior to sowing to prevent insect infection. Seeds were covered with clear plastic wrap and stratified at 4°C in the dark for at least 72 hours to break dormancy and coordinate germination. Stratified seeds were transferred to a temperature controlled Conviron S10H growth chamber (Conviron, Winnipeg, MB, Canada) for cultivation under standard growth conditions: 12-hour photoperiod (08:00-20:00), 100-150 μ mol photons m^-2^ s^-1^, 20 °C (± 2 °C), 55% (± 5 %) relative humidity. Upon germination, clear plastic wrap was slowly removed over 7-10 days, to maintain high humidity until seedlings were well-established and to avoid humidity shock. Plants were watered every 2-3 days depending on soil moisture, avoiding pooling of water to prevent algal and fungal growth. Each of three daily EL treatments consisted of 60 minutes of 1000 μ mol photons m^-2^ s^-1^ using a mixture of metal halide and high-pressure sodium lamps as previously described (Crisp *et al.* 2017; Ganguly *et al.* 2018).

For MethylC- and mRNA-seq experiments, developmentally equivalent leaves were harvested for comparisons (true leaves 4 - 9 in order of emergence). Harvested tissue was flash-frozen in liquid N_2_ and ground into a fine powder using a 1/8″ steel ball bearing, in a 1.5 ml Eppendorf tube, with 1 min shaking at 25 Hz in the *Tissue Lyser II* (Qiagen, Hilden, Germany). Ground tissue was stored at −80 °C. Approximately 20 and 30 mg of ground tissue was used for extracting total RNA and genomic DNA, respectively. MethylC-seq of WT and *strs2* was performed on paired tissue samples by using aliquots of ground frozen tissue from the same harvested plant.

### High-Throughput Phenotyping

For all genotypes the development of 10-day old (post-germination) Arabidopsis seedlings was followed for 20 days (until 4-weeks of age) by measuring plant area and rosette compactness. The *PlantScreen Compact System* (Photon Systems Instruments, Brno, Czech Republic), a high-throughput platform for digital plant phenotyping, was used to measure plant area and rosette compactness at 08:00 daily. Images were analysed using the *PSI RGB-IR Analyzer* software (version 1.0.0.2; Photon Systems Instruments, Brno, Czech Republic). Program settings were adjusted as needed to achieve well-defined plant areas and minimise background noise. The total number of macroscopically visible true leaves (cotyledons excluded) per rosette were also manually counted daily.

### Monitoring PSII performance using chlorophyll fluorescence measurements

PSII photochemistry was probed in vivo using measures of chlorophyll fluorescence (Baker 2008) using a PSI FluorCam (Photon System Instruments; Brno, Czech Republic). Measures were taken between 11:00-14:00 across the adaxial side of 30 minute dark adapted rosettes as performed previously (Ganguly *et al.* 2018). In this study, intermittent measures of chlorophyll fluorescence, specifically F_t_, F_m_, and F_m_′, were taken after a saturating pulse (3000 µ mol photons m^-2^ sec^-1^) across 10-minutes in actinic light (700 μ mol photons m^-2^ sec^-1^) followed by a 4-minute dark period (https://goo.gl/RuYDE2). The resulting fluorescence images were analysed using the FluorCam 7 software (v1.1.1.4; Photon System Instruments, Brno, Czech Republic). Chlorophyll fluorescence signals were analysed across whole rosettes. To better quantify light energy partitioning between photochemistry, light-regulated thermal dissipation, and other non-light induced quenching, such as chlorophyll fluorescence, we adopted the yield terms of ΦPSII, ΦNPQ, and ΦNO (Table 3), which sum to unity and do not require measures of F_o_ (Hendrickson *et al.* 2004; Kramer *et al.* 2004).

**Table 3.** Diagnostic PSII parameters and their biological relevance

| Parameter | Equation | Interpretation |
| --- | --- | --- |
| $\Phi_{PSII}$ | $1 - F_t/F_m'$ | Fraction of light absorbed by PSII for photochemistry. |
| $\Phi_{NPQ}$ | $F_t/F_m' - F_t/F_m$ | Fraction of light absorbed by PSII that is lost via thermal dissipation ($\Delta pH$- and xanthophyll-regulated processes). |
| $\Phi_{NPQ}$ | $F_t/F_m$ | Fraction of light absorbed by PSII that is lost via chlorophyll fluorescence. |

### MethylC sequencing

Genomic DNA was extracted from ground tissue using the *DNeasy Plant Mini Kit* (Qiagen, Netherlands) according to the manufacturer’s instructions, and quantified using a *ND-1000* Spectrophotometer. 70 ng of *Covaris* sheared gDNA (average fragment size = 200 bp) was bisulfite converted using the *EZ DNA Methylation-Gold Kit* (Zymo Research, CA, USA) according to the manufacturer’s instructions. Bisulfite converted DNA was used to create dual-indexed MethylC-seq libraries using the *Accel-NGS Methyl-Seq DNA Library Kit* paired with the *Accel-NGS Methyl-Seq Dual Indexing Kit* (Swift Biosciences, MI, USA) according to manufacturers’ instructions. All libraries were amplified in a 7-cycle indexing PCR reaction. All clean-ups were performed using either *AMPure XP beads* (Beckman Coulter, CA, USA) or *Sera-mag SpeedBeads* (GE Healthcare, Buckinghamshire, UK). A *LabChip GXII* (Perkin Elmer, MA, USA) was used to determine library molarity and fragment size distribution. Libraries were pooled in equimolar ratios and sequenced on one HiSeq2500 flow cell (100 bp single-end) at the *ACRF Biomolecular Research Facility* (Australian National University, ACT, Australia). In-depth details of library preparation are also described on protocols.io (https://goo.gl/vfwtEU).

Raw reads were quality controlled using *FastQC* (v0.11.2) with reads filtered and trimmed using *Cutadapt* (v1.9) and *Trim Galore!* (v.0.3.7) under default parameters. Single-end alignments of trimmed raw reads were aligned to the TAIR10 reference using *Bismark* (v0.14.5) (Krueger and Andrews 2011) and *Bowtie2* (v2.2.9) (Langmead and Salzberg 2012) with the flags -N 0 and –L 20. Per cytosine methylation levels were calculated using *Bismark methylation extractor* with default settings. Only cytosines with read depth > 3X were retained for further analysis. Bisulfite conversion efficiency was calculated as the proportion of methylated cytosines in the CHH context within the chloroplast genome, which itself should be fully unmethylated. Alignment metrics are provided in Supplementary Dataset 1. Weighted methylation levels were used to calculate the proportion of CG, CHG, and CHH methylation to account for sequencing depth (Schultz *et al.* 2012). This output was binned into 50 kbp regions (filtered for read depth > 15X) across the genome to construct chromosomal level metaplots of methylation levels using *BEDTools (Quinlan and Hall 2010)*.

Weighted methylation levels at single cytosines was utilized in DMR identification using *DSS* (v2.28.0) with default settings, including smoothing (*smoothing span* = 100) to improve methylation estimates (Feng *et al.* 2014; Wu *et al.* 2015). Differentially methylated cytosines (DMCs) were called (*DMLtest*) based on the posterior probabilities (q-value < 0.05) for a threshold in methylation difference (delta) at each cytosine in a context-specific manner: 0.5 CG, 0.2 CHG, and 0.1 CHH. Subsequently, DMRs are called based on adjacent statistically significant DMCs (*callDMR* with default parameters). These were refined by removing regions with a merged test statistic (*areaStat*) and estimated methylation difference in the lowest quartile, per sequence context, to remove DMRs containing DMCs of opposite direction. The final list of DMRs were assigned genomic positions, or overlapped between comparisons, using *BEDTools* and the *Araport11* annotation (Quinlan and Hall 2010; Cheng *et al.* 2017). Code used for analyses are available on *Github* (https://goo.gl/wsQrJT).

Independent WT and *drm1drm2* MethylC-seq profiling was accessed from <u>GSE39901</u> and <u>GSE38286</u>. DRM1/2-dependent RdDM sites were determined as the CHH hypo-DMRs identified in common between *drm1drm2* and three independent WT samples using *DSS* with same parameters used herein (Stroud *et al.* 2015).

### mRNA sequencing

Total RNA was extracted with *TRIzol* (Life Technologies) using an adapted protocol (Allen *et al.* 2010). Briefly, ground tissue was lysed in 1 ml *TRIzol* and mixed by gentle inversion and incubated at room temperature for 5 minutes. Subsequently, 200 μl chloroform was added and shaken vigorously to mix. Samples were centrifuged at 14,000 rcf for 10 min at 4°C to separate the resulting upper aqueous phase from the organic phase. Chloroform extraction was repeated twice, transferring 400-600 μl then 300-400 μl aqueous phase to a new microfuge tube following each extraction. RNA was precipitated by adding an equal volume of 100% isopropanol and mixing by inversion before incubating at −20°C overnight. RNA was recovered by 4°C centrifugation at 20,000 rcf for 20 min, the supernatant discarded, and the pellet washed with 80% ethanol and centrifuged at 7,500 rcf for a further 3 min. The supernatant was discarded and the pellet air-dried prior to resuspension in 50 μl DEPC-treated H_2_O. All purified RNA was stored at −80°C. RNA quantity was assessed using *ND-1000 spectrophotometer* (NanoDrop Technologies). RNA quality was assessed using the *LabChip GXII* (Perkin-Elmer) for RIN > 6.5.

Poly(A)-enriched RNA-sequencing (mRNA-seq) libraries were prepared using the *Illumina TruSeq Stranded mRNA Sample Preparation Kit* (Illumina, CA, USA) using 1.3 μg input of extracted total RNA. The following modifications to the manufacturer’s instructions were made: reagent volumes were adjusted for 1/3 reactions; and Invitrogen SuperScript III Reverse Transcriptase (Invitrogen, Life Technologies Australia Pty Ltd) was used for first strand synthesis with adjusted reaction temperature of 50 °C. Libraries were constructed using *Illumina TruSeq RNA Single Indexes* (Set A and B; Illumina, CA, USA) in a 14-cycle indexing PCR reaction; all clean-ups were performed using *RNAClean XP beads* (Beckman Coulter, CA, USA). A *LabChip GXII* (Perkin Elmer, MA, USA) was used to determine library concentration and fragment size distribution, using a DNA High Sensitivity Kit. mRNA-seq libraries were pooled in equal molar ratios and sequenced (75 bp single-end) on a *NextSeq500* at the ACRF Biomolecular Research Facility (Australian National University, ACT, Australia).

Raw reads were diagnosed using *FastQC* (v0.11.2). Due to strong nucleotide sequence content bias *Trim Galore!* and *Cutadapt* were used to trim low-quality reads with PHRED score < 20 (-q 20) and to make a hard clip of 10 bp and 1 bp from the 5’ and 3’ ends, respectively. Single-end alignments of trimmed raw reads were aligned to the *TAIR10* reference genome using *Subread* (v1.6.2) (Liao *et al.* 2013) with the flags -t 0 and -u to report uniquely mapping reads, prior to sorting, indexing and compressing using *Samtools* (v1.2). Alignment metrics are provided Supplementary Dataset 3. Transcript quantification was performed at the gene-level with the *Araport11* annotation (Cheng *et al.* 2017) using *featureCounts* (with flag -s 2 for reverse strand specificity).

Differential gene expression analyses were performed using the *edgeR* quasi-likelihood pipeline (Chen *et al.* 2016). Reads mapping to ribosomal RNA and organellar transcripts were removed; only loci containing counts per million (CPM) > 1 in at least three samples were examined. After this filtering, 17,657 loci were retained for analysis. The trimmed mean of M-values (TMM) method was used to normalise transcript abundance between libraries to account for sequencing depth and composition. Subsequently, generalised linear models were fitted to estimate dispersion (*glmQLFit*) allowing for differential expression testing, employing quasi-likelihood F-tests (*glmQLTest*) and controlling for false discovery rates due to multiple hypothesis testing (FDR adjusted p-value < 0.05). Gene ontology enrichments were examined using the *statistical overrepresentation test* (Binomial test with FDR correction) from the *PANTHER* classification suite (Mi *et al.* 2013). Code used for RNA-seq analyses are available on *Github* (https://goo.gl/b7x5rc).

### Statistical analyses

Statistical analyses and data visualisation was performed using *R* (v 3.5.0) with the *tidyverse* package (v 1.2.1). Linear mixed-effects models were fitted using the *lme4* package (v 1.1-17) (Bates *et al.* 2015) to account for both fixed (e.g. genotype, condition, time) and random effects (e.g. experimental design, blocking factors). Model fit was assessed using the conditional R^2^ value calculated using the *piecewiseSEM* package (v 2.0.2) (Lefcheck 2016). Fitted models allowed estimation of marginal means and 95% confidence intervals using the *emmeans* package including *post hoc* contrasts between factors with FDR p-value correction (v 1.2.1). All analyses were performed on single data points, representing individual biological replicates (independent plants). Hypergeometric testing was performed using the *phyper* function to test for significant overlaps, taking into account the 17,657 detected transcripts and the number of stress signalling-associated genes overlapped.

## Data accessibility

All sequencing data generated in this study has been deposited at the NCBI GEO repository: <u>GSE121150</u>.

## Supplemental Datasets

Supplementary Dataset 1 MethylC sequencing statistics

Supplementary Dataset 2 *strs2* DMRs

Supplementary Dataset 3 mRNA sequencing statistics

Supplementary Dataset 4 Differentially expressed genes following triggering stress in naive plants

Supplementary Dataset 5 Differentially expressed genes following triggering stress in primed plants

Supplementary Dataset 6 Dampened transcripts

Supplementary Dataset 7 Enriched GO terms among dampened transcripts

Supplementary Dataset 8 Genotyping primers

## Acknowledgements

We would like to acknowledge the advice received from Prof. Wah Soon Chow on chlorophyll fluorescence based measures of photosynthetic performance. We would also like to acknowledge the Biomolecular Resource Facility at the ANU for performing Illumina sequencing, and Dr Terry Neeman for advice on statistical analyses. This project was also supported with the provision of plant growth facilities by the Australian Plant Phenomics Facility and computational infrastructure by the National Computational Infrastructure, both supported under the National Collaborative Research Infrastructure Strategy of the Australian Government.

This project was funded by the Australian Research Council Centre of Excellence in Plant Energy Biology (CE140100008). DRG was supported by the Grains Research and Development Council (GRS10683) and an Australian Research Training Program (RTP) Scholarship. SRE was funded by a Discovery Early Career Researcher Award (DE150101206).

The authors declare no conflicts of interest.

---

**Figure S1.**
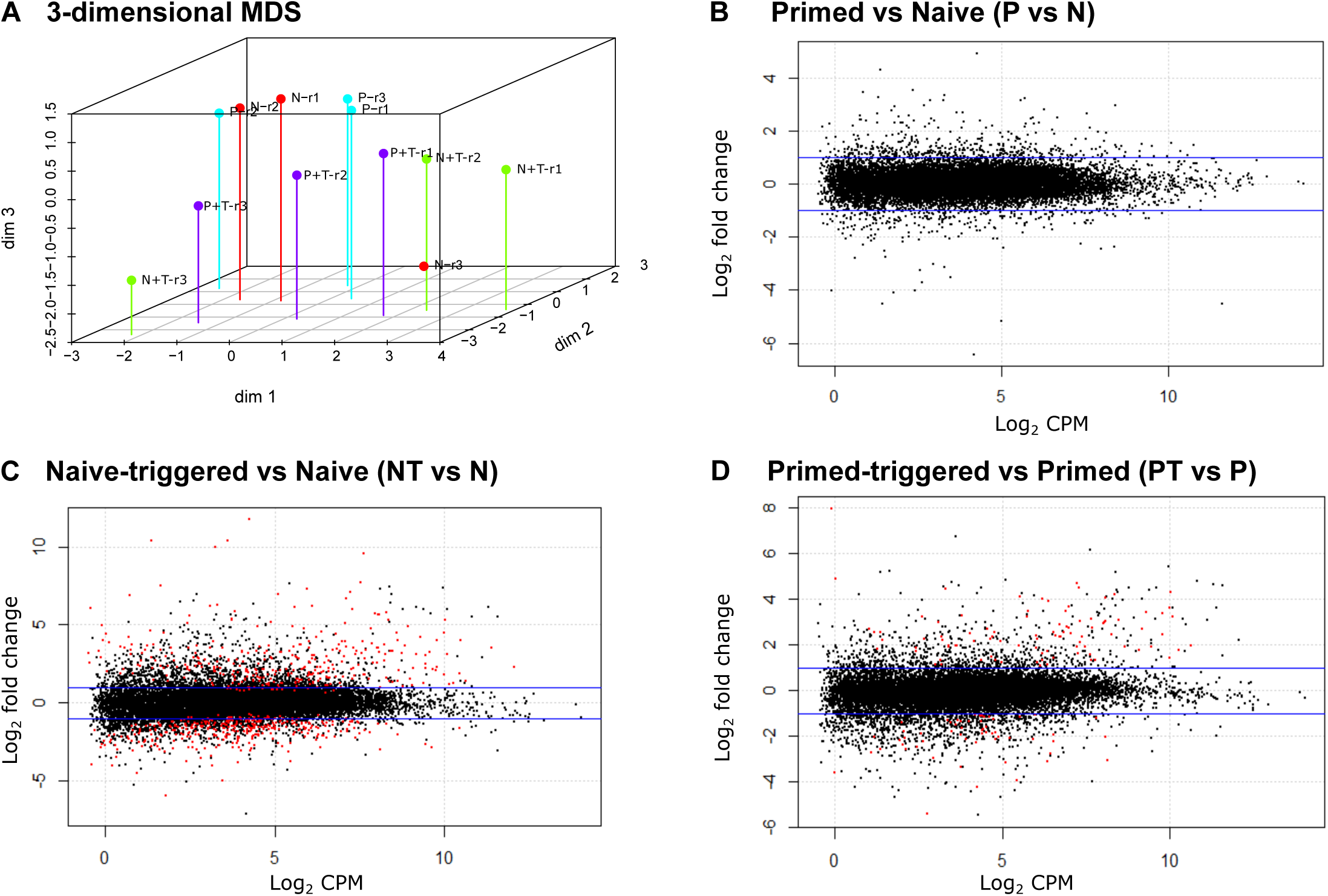
Exploratory transcriptome analysis of naive and primed plants. **A** Multi-dimensional scaling plot of all samples where distance reflects the typical log_2_ fold change between samples. **B-D** Mean-difference plots for **B** primed versus naive libraries, **C** naive-triggered versus naive libraries, and **D** primed-triggered versus primed libraries with smearing of low abundance transcripts. Each dot represents a transcript plotted by its average abundance (log_2_ CPM) against the log_2_ fold change from the specified comparison. Red dots indicate differentially expressed transcripts as determined by quasi-likelihood F-tests (FDR < 0.05). Blue lines denote 2-fold change.

## Notes

#### Summary of Updates

Updated authors and substantive changes to text.

https://www.ncbi.nlm.nih.gov/geo/query/acc.cgi?acc=GSE121150

